# A network-based approach reveals the dysregulated transcriptional regulation in non-alcohol fatty liver disease

**DOI:** 10.1101/2021.07.24.453650

**Authors:** Hong Yang, Muhammad Arif, Meng Yuan, Xiangyu Li, Ko Eun Shong, Hasan Turkez, Jens Nielsen, Mathias Uhlén, Jan Borén, Zhang Cheng, Adil Mardinoglu

## Abstract

Non-alcohol-related fatty liver disease (NAFLD) is a leading cause of chronic liver disease worldwide. We performed network analysis to investigate the dysregulated biological processes in the disease progression and revealed the molecular mechanism underlying NAFLD. Based on network analysis, we identified a highly conserved disease-associated gene module across three different NAFLD cohorts and highlighted the predominant role of key transcriptional regulators associated with lipid and cholesterol metabolism. In addition, we revealed the detailed metabolic differences between heterogenous NAFLD patients through integrative systems analysis of transcriptomic data and liver-specific genome-scale metabolic model. Furthermore, we identified transcription factors (TFs), including SREBF2, HNF4A, SREBF1, YY1 and KLF13, showing regulation of hepatic expression of genes in the NAFLD-associated modules and validated the TFs using data generated from a mouse NAFLD model. In conclusion, our integrative analysis facilitated our understanding of the regulatory mechanism of these perturbed TFs and associated biological processes.

## INTRODUCTION

Non-alcohol fatty liver disease (NAFLD) is considered as one of the most important causes of liver disease, worldwide (Asrani et al., 2019). The global prevalence of NAFLD was estimated to be 25% and has increased rapidly (Huang et al., 2021; Younossi et al., 2018; Younossi et al., 2016). NAFLD is characterized by the hepatic accumulation of triglycerides, spanning from simple non-alcohol fatty liver (NAFL) to non-alcohol steatohepatitis (NASH) that might progress to cirrhosis and hepatocellular carcinoma (HCC) (Friedman et al., 2018; Huang *et al*., 2021; Ioannou et al., 2019). Moreover, NAFLD is strongly associated with obesity, diabetes and cardiovascular disease; therefore, it drastically increased in these patient groups (Golabi et al., 2019; Ye et al., 2020; Younossi et al., 2019). However, no effective therapies are yet approved for NAFLD, even though extensive research activities have been carried out (El-Agroudy et al., 2019; Mullard, 2020; Newsome et al., 2021; Stower, 2021). Hence, a comprehensive understanding of the underlying molecular mechanism of NAFLD is critical for the development of novel approaches for its prevention and treatment.

Biological networks provide a robust framework for integrating omics data, elucidating pathophysiological responses and revealing the underlying molecular mechanisms involved in the progression of disease (Calabrese et al., 2017; Mardinoglu et al., 2018; Nayak et al., 2009). Biological networks, including protein-protein interaction networks (PPINs), transcriptional regulatory networks (RNs), gene co-expression networks (GCNs), genome-scale metabolic models (GEMs) and integrated networks (INs), are widely used in systems analysis (Mardinoglu *et al*., 2018). The central goal of biological network analysis is to identify critical functional units (so-called modules) and their constituent genes (Califano et al., 2012; Choobdar et al., 2019). Such functional modules often linked to disease processes, in which key drivers are highly enriched and provide insights into the disease pathogenesis (Cerami et al., 2010; Huan et al., 2013; Wainberg et al., 2021). In particular, GCNs for 17 human cancers and 46 human tissues have been generated and used to gain insights into disease mechanisms by identifying the key biological components of the cancers or tissues (Arif et al., 2021; Lee et al., 2018; Uhlen et al., 2017). GEMs, reconstructed by incorporating all biochemical reactions and transport processes in a cell or tissue, have been extensively used to discover potential biomarkers and drug targets, as well as to reveal the mode of action of a drug (Lewis and Kemp, 2021; Mardinoglu *et al*., 2018).

To date, GCNs have been used for investigating the causal mechanisms underlying NAFLD using mouse population data (Chella Krishnan et al., 2018) and human population data (Zhang et al., 2020) and for integrative analysis of mouse model data and patients data (Saeed, 2021). However, there is still a lack of holistic studies in which adequately samples covering a large spectrum of disease severity were analysed. Moreover, the heterogeneity of clinical manifestations among NAFLD patients acts as an essential impediment for the discovery of critical pathogenic drivers, and thus systematic analysis is needed (Alonso et al., 2017). Recently, several liver-biopsy proved transcriptomics data from large patients’ cohorts had been conducted (Azzu et al., 2021; Govaere et al., 2020; Hoang et al., 2019), and these datasets may be used to provide significant functional insights based on network analysis that cannot be derived from individual gene-level analysis.

In this study, we employed an integrative systems biology approach by integrating NAFLD transcriptomics data with biological networks and elucidated the molecular mechanisms underlying NAFLD progression. We first generated GCNs for liver tissue of normal and NAFLD patients based on transcriptomics data and identified the perturbated modules associated with the severity of NAFLD. Second, we employed a liver-specific GEM called *iHepatocytes2322* to analyse the differential expression data, and gained insights into detailed metabolic differences in NAFLD. Third, we subsequently validated the perturbated modules using transcriptomics data from another two independent studies and highlighted the disease-associated modules that are conserved across multiple NAFLD cohorts by combing functional and topological similarities. Next, we used a liver cancer dataset in the cancer genome atlas (TCGA) to investigate if the dysregulated expression of genes in the disease-associated modules is relevant to patient outcome. Finally, we performed transcription factor (TF)-target regulatory network analysis, identified TFs that regulate disease-associated modules and validated those TFs using the transcriptomics data from a mouse NAFLD model fed by high sucrose diet (HSD).

## RESULTS

### Generation of co-expression networks for liver tissue

We identified the robust co-expressed genes showing transcriptional differences between the liver tissue of normal subjects and NAFLD patients. We first constructed GCNs of liver tissue based on the transcriptomics data from the Genotype-Tissue Expression (GTEx) database (GTEx analysis V8) (Consortium, 2013) and NAFLD cohort including 10 normal samples, 50 patients with NAFL and 155 patients with NASH (Govaere *et al*., 2020) (Figure 1A). We filtered out lowly-expressed genes for each dataset based on their mean gene expression level (TPMs <1) and performed Spearman’s rank correlation test between each gene pair. All p values were adjusted by FDR correction (Benjamini-Hochberg). Afterwards, we remained gene pairs with significantly positive correlation (coefficient > 0 with FDR < 0.05) on the networks (Figure 1A) and used the Leiden algorithm (Traag et al., 2019) to identify modules of genes from the network. In total, Leiden graph-based clustering identified six and five modules of genes in the GTEx cohort and NAFLD cohort, respectively. Each module in the same cohort consists of uniquely assigned genes with a substantial similarity between gene expression (Figure 1B; Dataset S1). Of note, we found that gene members of any module in the GTEx cohort were different from that of modules in the NAFLD cohort even though 95.9% genes comprising modules in the NAFLD cohort were included by GTEx modules (Figure 1C).

**Figure 1.**
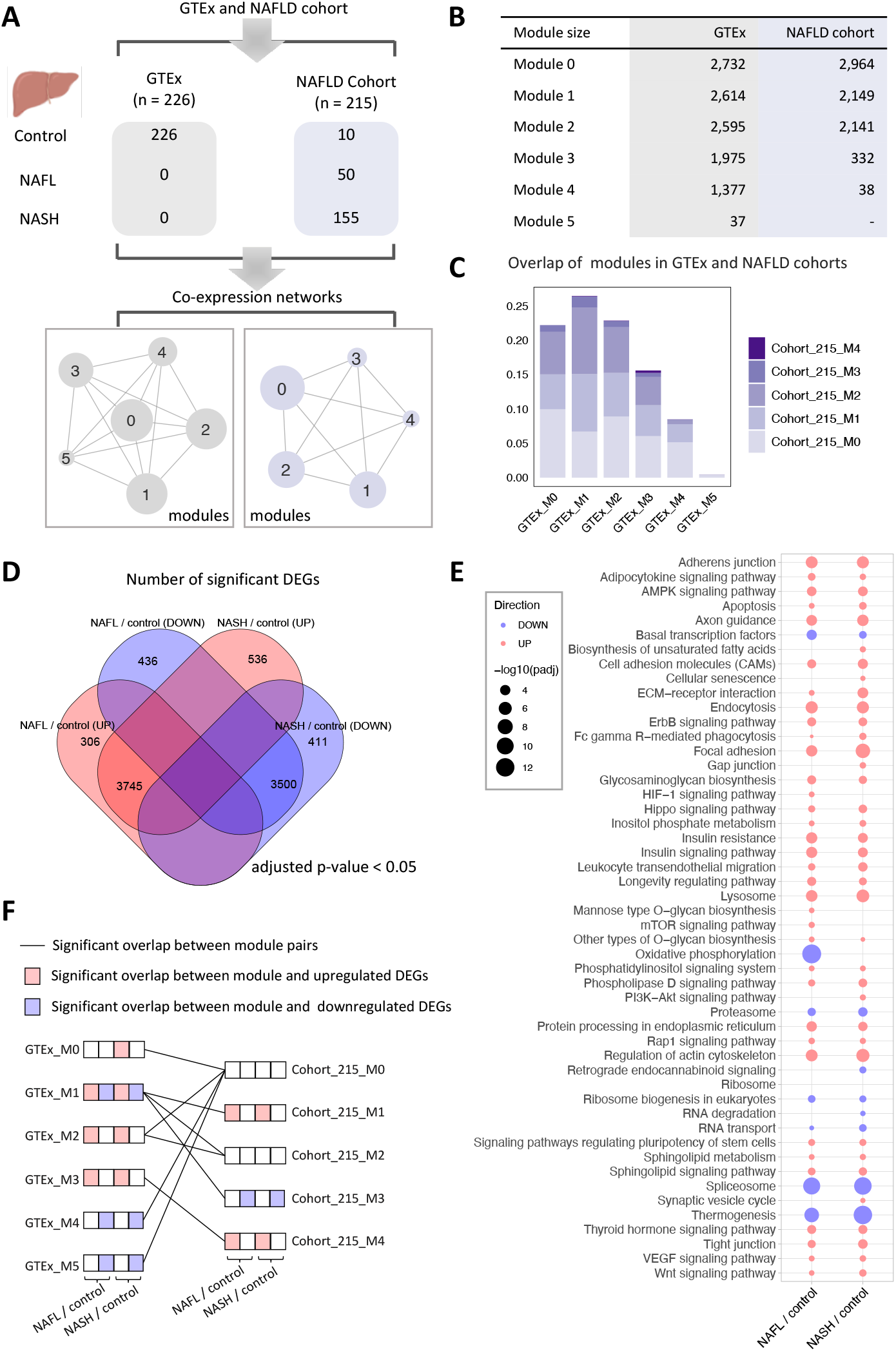
Sample information of studied cohorts and construction of co-expression networks. (**A**) Transcriptome data of liver tissue were obtained from GTEx, NAFLD cohort with 226 and 215 samples ranging from normal, NAFL, NASH, respectively. Spearman rank-order correlation coefficient analysis was applied to calculate the correlation between gene pairs after removing the lowly expressed genes (TPMs <1), and the Leiden algorithm was used to detect modules of significantly correlated genes. The label (number) of the module assigned by the algorithm. (**B**) The numbers of genes consist of the individual module in each cohort. (**C**) The bar plot shows the proportions of genes in each module identified in the NAFLD cohort compared to each module specified in the GTEx cohort. (**D**) The Venn diagram of differentially expressed genes (adjusted p-value < 0.05) between patients with NAFLD and control samples. (**E**) KEGG pathway analysis showing pathways that were significantly altered in patients with NAFLD. Up-regulated and downregulated pathways are shown in blue and red, respectively— only pathways with adjusted p-value (padj) < 0.01 are presented (see also Dataset S3. The size of the bubble is scaled by -log_10_(padj) for each KEGG pathway term. (**F**) Significant (p < 0.05, hypergeometric test) overlap between module pairs between GTEx and NAFLD cohorts and overlap between the module and dysregulated genes associated with NAFLD. GTEx, genotype-tissue expression; NAFLD, non-alcohol fatty acid disease; NAFL, non-alcohol fatty liver; NASH, non-alcohol steatohepatitis; TPMs, transcripts per kilobase million.

### Identification of perturbated modules in NAFLD

We investigated whether the differences in module composition correlated with the molecular changes underlying NAFLD progression. We first identified differentially expressed genes (DEGs) to reveal the global transcriptomic differences in the liver of patients with NAFLD. We observed that 4,051 and 4,281 genes were significantly upregulated (adjusted p-value < 0.05) between NAFL and control samples and between NASH and normal samples, respectively (Figure 1D; Dataset S2). Enrichment analysis in KEGG pathway showed that the upregulated DEGs are mostly enriched in the pathways associated with endocytosis, axon guidance, adherens junction, insulin resistance and insulin signalling (Figure 1E; Dataset S3). Moreover, we found that 3,936 and 3,911 genes were significantly downregulated between NAFL and control samples and between NASH and normal samples, respectively (Figure 1D; Dataset S2). Enrichment analysis showed that downregulated DEGs enriched in pathways associated with oxidative phosphorylation, spliceosome, thermogenesis, and proteasome (Figure 1E).

We also examined the enrichment of those dysregulated DEGs associated with NAFLD in each co-expression module identified in GTEx and NAFLD cohort data. The results showed that module 1 and module 3 of the NAFLD cohort with 215 samples (cohort_215_M1 and cohort_215_M3) are significantly enriched (hypergeometric test p-value ≈ 0) by upregulated and downregulated DEGs associated with NAFLD, respectively (Figure 1F). In particular, 66.6% of genes (1,431 out of 2,149) in cohort_215_M1 and 94.6% of genes (314 out of 332) in cohort_215_M3 are significantly upregulated and downregulated in NAFL vs control groups, respectively. 65.8% of genes in cohort1_215_M1 and 94% of genes (312 out of 332) in cohort_215_M3 are significantly upregulated and downregulated in NASH vs control groups, respectively. Notably, we found both cohort_215_M1 and cohort_215_M3 are significantly overlapped (hypergeometric test p-value = 2.17×10^-14^ and 3.11×10^-8^) with module 1 in GTEx cohort (GTEx_M1), which were overrepresented by both upregulated and downregulated DEGs associated with NAFLD (Figure 1F). Interestingly, KEGG enrichment analysis of genes in those modules suggests that the significantly enriched pathways are consistent with the dysregulated pathways enriched by DEGs (Figure 1E; Figure S2A). Moreover, we found cohort_215_M4 are significantly enriched by upregulated genes (19 and 25 out of 38 in NAFL vs control and NASH vs control, respectively) and only significantly overlapped with GTEx_M3 (hypergeometric test p-value = 4.11×10^-13^). Taken together, co-expression network analysis identified modules of genes that are significantly perturbated in patients with NAFLD.

### Altered metabolism in NAFLD patients

To further evaluate the detailed metabolic changes underlying NAFLD progression, we identified reporter metabolites (Patil and Nielsen, 2005), around which the most significant transcriptional changes occur, using differential expression data from NAFLD and network topology provided by *iHepatocytes2322* (Mardinoglu et al., 2014). (Figure S2B). Such reporter metabolites can thus be used to identify the key dysregulated regions of the metabolic network. A total of 321 metabolites were significantly (p-value <0.05) associated with upregulated genes in either NAFL vs control or NASH vs control (Figure 2; Figure S2C; Dataset S4), Among these, the most significant reporter metabolites associated with upregulated genes in NAFL vs control were those involved in arginine and proline metabolism, glycerophospholipid metabolism, and nucleotide metabolism. The top reporter metabolites associated with upregulated genes between NASH and control samples involved in beta-oxidation of fatty acids, cholesterol biosynthesis, and chondroitin/heparan sulphate biosynthesis. Chondroitin sulphate (CS) and heparan sulphate (HS) are the essential components of proteoglycans (PG), which have been proposed as potential biomarkers for NASH diagnosis and staging of NAFLD by integrative analysis of transcriptomic data obtained from patients with NAFLD and GEM (Mardinoglu *et al*., 2014). The analyses from the current investigation utterly consistent with the previous study. In addition, we observed 215 metabolites were significantly associated with downregulated genes in NAFLD, involving in folate metabolism and oxidative phosphorylation (Figure 2; Figure S2D; Dataset S4).

**Figure 2.**
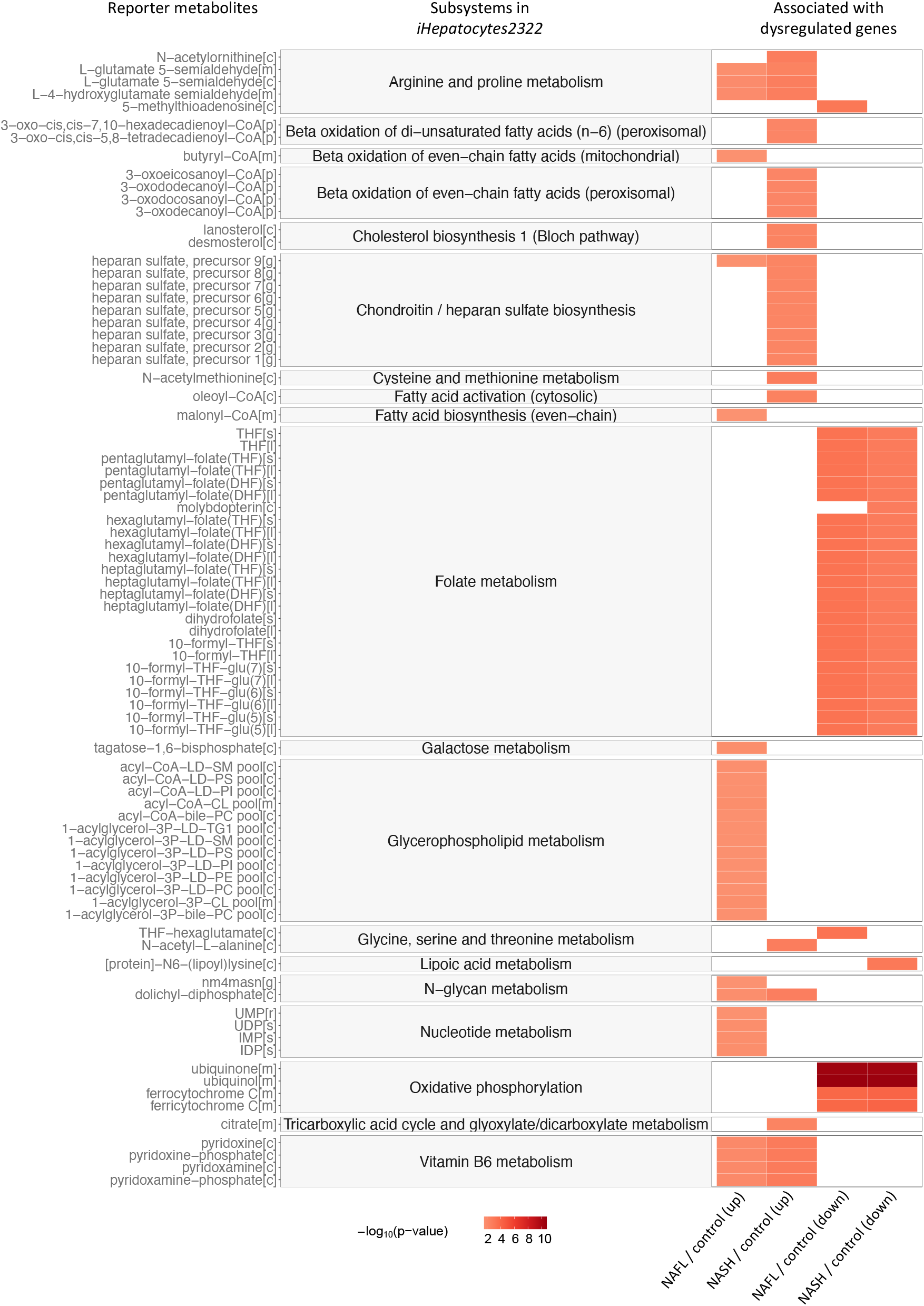
The most significant reporter metabolites between patients with NAFLD and control samples through the employment of *iHepatocytes2322*. Reporter metabolites were calculated for the upregulated and downregulated genes for each comparison. Top30-ranked reporter metabolites and subsystems in *iHepatocytes2322* associated with up-regulated and down-regulated genes in each comparison are presented, respectively. Colour is proportional to the minus logarithm of the p-value (-log_10_(p-value)), see also Dataset S4.

### Validation of perturbated modules in two independent NAFLD cohorts

To validate whether modules related to significant transcriptomics and metabolic changes in patients with NAFLD can truly reflect the perturbations in a disease-specific manner, we analysed GCNs generated using liver-biopsy proved transcriptomics datasets from two independent NAFLD cohorts with 75 and 58 samples, respectively (Azzu *et al*., 2021; Hoang *et al*., 2019) (to avoid repeated IDs, we assigned cohort 1 to the studied NAFLD cohort, and 2 and 3 for NAFLD cohorts for validation in the downstream analysis). By the same method constructing GCN for the first NAFLD cohort as described in Method, we identified four and eight modules of genes in NAFLD cohort 2 and 3, respectively (Figure S3A&B). To explore module similarity among NAFLD cohorts, we calculated the Jaccard index between each pair of modules from different NAFLD cohorts and performed hypergeometric test to evaluate the significance of the observed overlap in gene members (Figure 3A, B&C; Dataset S1). To begin with, we tested the modules between NAFLD cohort 1 and cohort 2. The results showed that genes in cohort1_215_M4 were only significantly overlapped (29 out of 38; Jaccard index = 0.388; hypergeometric test p-value = 1.66×10^-69^, Figure 3A&D) with genes in module 3 of NAFLD cohort 2 with 75 samples (cohort2_75_M3). We next tested the module pairs in NAFLD cohort 1 and cohort 3, found cohort1_215_M4 also shared 29 genes with module 7 of NAFLD cohort 3 with 58 samples (cohort_3_58_M7) (Jaccard index = 0.434; hypergeometric test p-value = 3.73×10^-71^, Figure 3B&D). Interestingly, the genes in cohort2_75_M3 were significantly overlapped (45 out of 80; Jaccard index = 0.425; hypergeometric test p-value = 1.82×10^-86^, Figure 3C&D) with genes in cohort3_58_M7 as well.

**Figure 3.**
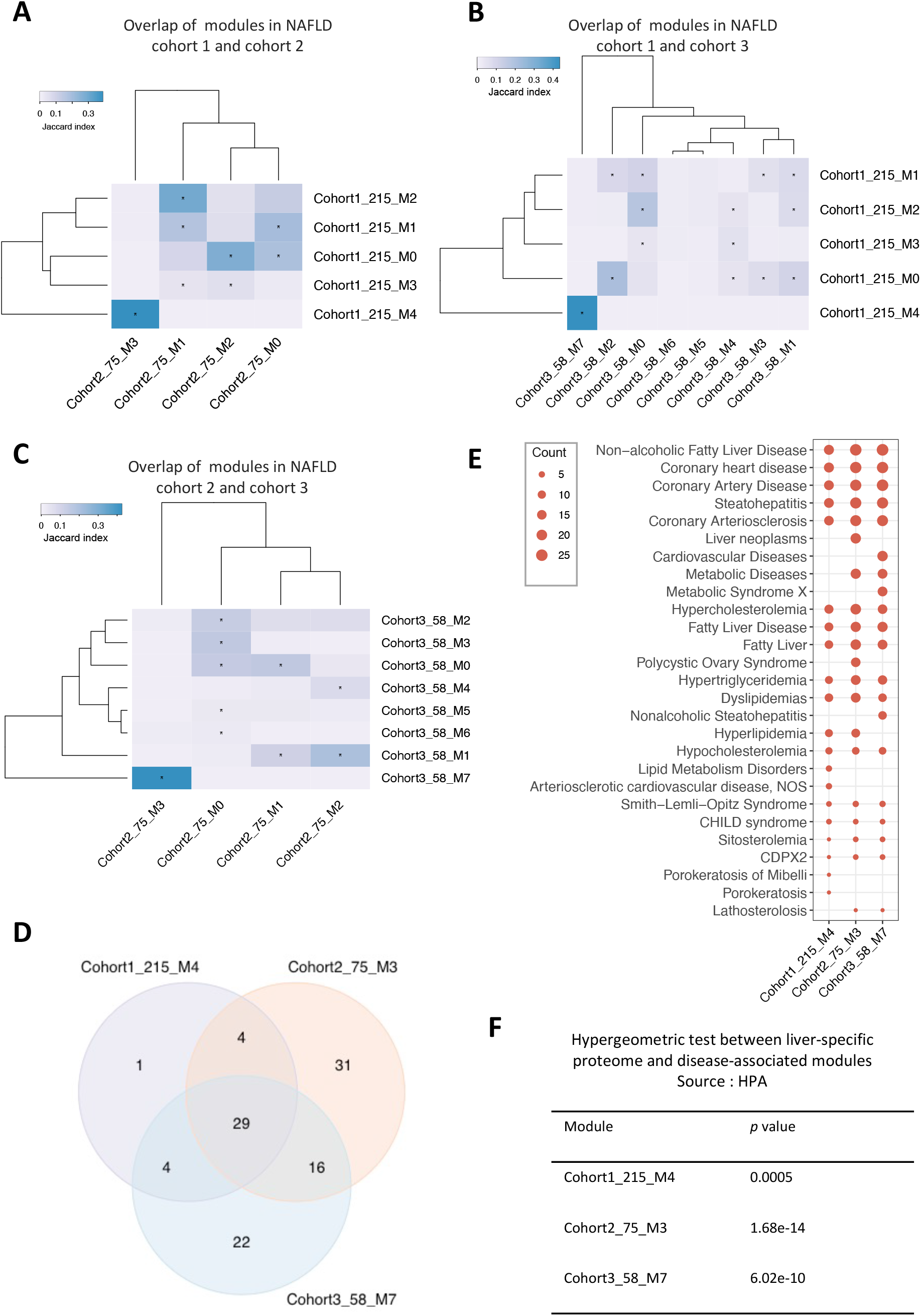
Validation of disease-related modules using two independent NAFLD cohorts. (**A-C**) Hierarchical clustering of Jaccard Index between module pairs from NAFLD cohort 1 and 2; NAFLD cohort 1 and 3; NAFLD cohort 2 and 3. Color scales representing the range of the Jaccard index. asterisk indicates the statistical significance of the overlap between gene members in any two modules from the different cohorts. (**D**) Venn diagram shows numbers of genes overlapped between cohort1_215_M4, cohort2_75_M3, and cohort3_58_M7. (**E**) Dot-plot heatmap shows top 20 significantly (‘q-value FDR B&H’ < 0.05) enriched diseases by genes in each module (cohort1_215_M4, cohort2_75_M3, and cohort3_58_M7). The size of each dot is proportional to the number of genes enriched in each disease term. (**F**) The table shows the results from a hypergeometric test between liver-specific proteome (HPA) and disease-associated modules in NAFLD cohorts, p-value less than 0.05.

We subsequently assessed the module similarity and overlap between any two modules between the GTEx cohort and NAFLD cohorts to validate if those conserved modules in NAFLD cohorts have a similar expression pattern in normal liver tissue. Hierarchical clustering of the Jaccard index between module pairs showed a distinct cluster consisting of cohort1_215_M4, cohort2_75_M3, and cohort3_58_M7 was only significantly over-represented by module 3 in GTEx cohort (GTEx_M3), which contains 1,975 genes (Figure 1B; Figure S3C). We found that more than 70% genes in cohort1_215_M4 (27 out of 38; hypergeometric test p-value = 4.11×10^-13^), cohort2_75_M3 (57 out of 80; hypergeometric test p-value = 4.97×10^-27^), and cohort3_58_M7 (50 out of 71; hypergeometric test p-value = 2.92×10^-24^) were included by module GTEx_M3. (Figure 4A).

**Figure 4.**
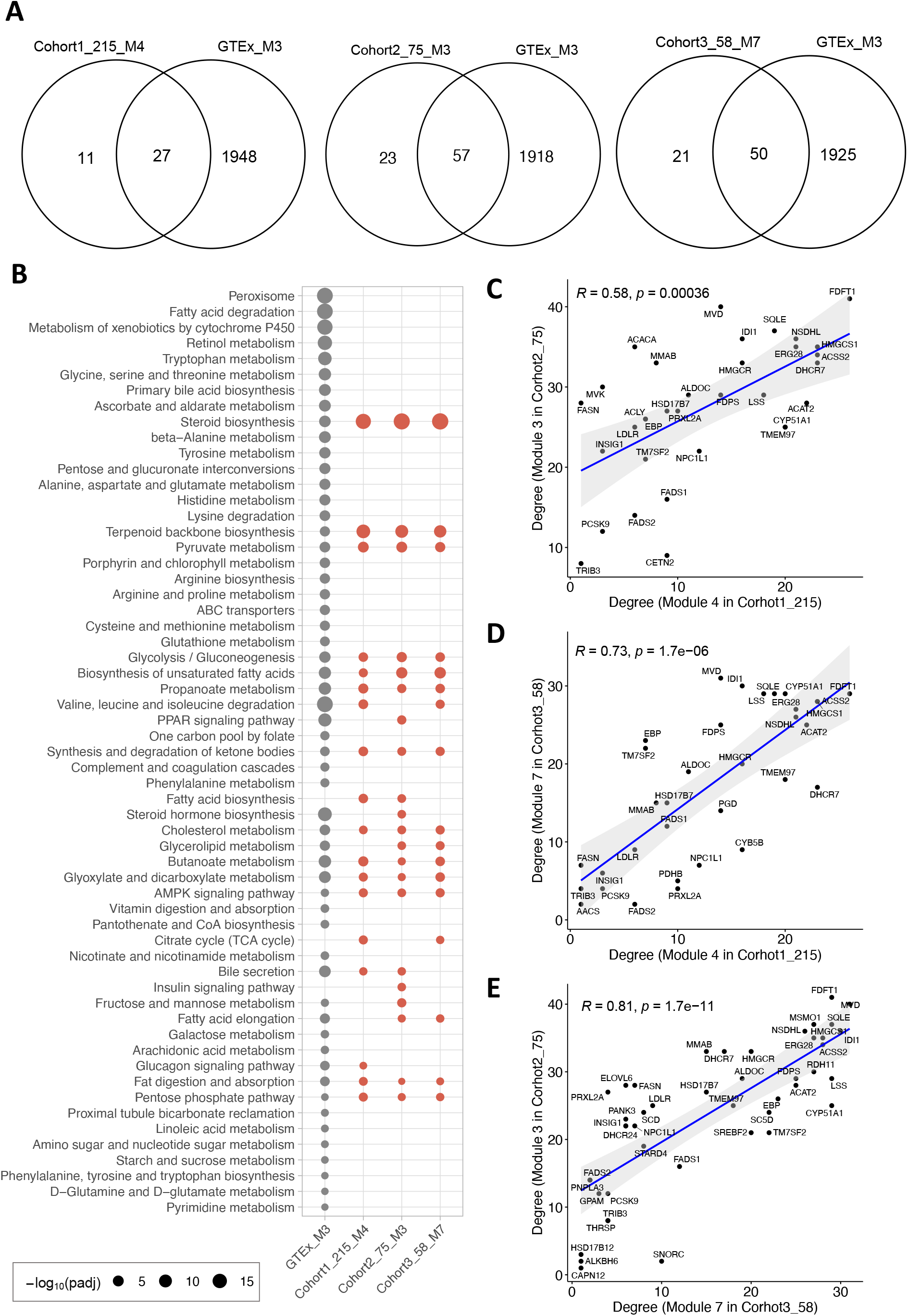
(**A**) Venn diagram shows numbers of genes overlapped between GTEx_M3 cohort and cohort1_215_M4, cohort2_75_M3, and cohort3_58_M7, respectively. (**B**) Dot-plot heatmaps are showing KEGG pathways enriched in different modules. The colour differences of dots indicate the studied cohort (GTEx or NAFLD) in which the module detected. The size of each dot is proportional to the significance (-log_10_(padj); padj represents ‘q-value FDR B&H’ with value < 0.05) of enrichment for each KEGG pathway term. (**C-E**) Correlation between degree among disease-associated modules from different cohorts. The correlation was evaluated by Spearman correlation coefficients.

For a systematic evaluation on biological functions related to the modules, we quantified the statistical significance of enrichment of genes with the association in disease-related gene sets obtained from DisGeNET database (https://www.disgenet.org/) (Pinero et al., 2020), liver-specific proteome in Human Protein Atlas (HPA) database (http://www.proteinatlas.org/) (Uhlen et al., 2015), and KEGG pathway gene sets. We found that genes in cohort1_215_M4, cohort2_75_M3, and cohort3_58_M7 were significantly over-represented by multiple liver disease-related gene sets, including NAFLD and steatohepatitis (Figure 3E; Dataset S5). Interestingly, these three modules were also significantly enriched in heart diseases, for instance, coronary heart disease, coronary artery disease, and coronary arteriosclerosis. We further evaluated the overlap of genes in each of three disease-associated modules with 936 liver-specific genes defined by HPA (Fagerberg et al., 2014; Uhlen *et al*., 2015; Yu et al., 2015). The results showed that genes in cohort1_215_M4 (10 out of 38; hypergeometric test p-value

= 0.0005), cohort2_75_M3 (30 out of 80; hypergeometric test p-value = 1.68×10^-14^), and cohort3_58_M7 (22 out of 71; hypergeometric test p-value = 6.02×10^-10^) are highly enriched with liver-specific genes (Figure 3F). In addition, we observed that genes in GTEx_M3, which shows high module similarity with those three modules identified in diseases cohorts, significantly enriched in the peroxisome, branched-chain amino acids (BCAAs; valine, leucine and isoleucine) degradation, and fatty acid degradation (Figure 4B). However, steroid biosynthesis and terpenoid backbone biosynthesis were the most significantly enriched pathways in all the three modules of disease cohorts. Moreover, fatty acid biosynthesis, citrate cycle (TCA cycle), and insulin signalling pathway were only significantly enriched in the modules of disease cohort(s).

### Topological features of genes in NAFLD-associated modules

The analysis of topological properties can provide important information about hub genes or other influential genes that significantly impact the dynamic of the module. To understand the interplay of genes in the module, we then obtained several key network properties using the “NetworkAnalyzer” in Cytoscape to analyse the disease-associated modules. In our workflow, we used degree and closeness centrality to evaluate the importance of nodes in a module. In an unredirected network, the degree of a node is the number of edges linked to this node and a node with a high degree has been considered as functionally significant (Doncheva et al., 2012). Genes with high closeness centrality are considered as the controlling point of molecular communication (Miryala and Ramaiah, 2019).

Topological analysis showed that gene *FDFT1* has the highest degree in both cohort1_215_M4 (26 edges) and cohort2_75_M3 (41 edges), whereas gene *MVD* has the highest degree in cohort3_58_M7 (Figure 4C-E; Figure S4; Figure S5A; Dataset S6). The other genes with high connectivity are *HMGCS1*, *DHCR7* and *ACSS2* (23 edges, respectively), and *ACAT2* (22 edges) in cohort1_215_M4; *MVD* (40 edges), *MSMO1* and *SQLE* (37 edges, respectively), and *IDI1* and *NSDHL* (36 edges, respectively) in cohort2_75_M3; *IDI1*(30 edges), *LSS*, *CYP51A1*, *FDFT1* and *SQLE* (29 edges, respectively) in cohort3_58_M7. Interestingly, we observed a highly positive correlation between the degree of 33 genes overlapped in cohort1_215_M4 and cohort2_75_M3 (Spearman’s correlation = 0.58, p = 0.00036, Figure 4C), which indicates that those two modules have a similar topological structure. Similarly, a highly positive correlation between degree of 33 genes shared by cohort1_215_M4 and cohort3_58_M7 (Spearman’s correlation = 0.73; p = 1.7×10^-6^, Figure 4D) and that of degree of 45 genes shared by cohort2_75_M3 and cohort3_58_M7 (Spearman’s correlation = 0.81; p = 1.7×10^-11^, Figure 4E) were also observed. Moreover, the top five genes with the highest closeness centrality in cohort1_215_M4, cohort2_75_M3 and cohort3_58_M7 are also highly conserved (Figure S5B). We also observed a strong correlation between closeness centrality of shared genes in any disease-associated module pairs of NAFLD cohorts (Figure S5C).

### Validation of topological features in an HCC cohort

Given that NAFLD has emerged as the fastest-growing cause of HCC (Huang *et al*., 2021; Ray, 2018). We next investigated whether the expression of the genes in NAFLD disease-associated modules, especially genes with high-connectivity, is predictive of patients with HCC using the Liver Hepatocellular Carcinoma dataset (TCGA-LIHC; https://portal.gdc.cancer.gov/projects/TCGA-LIHC) (Figure S6A). The results showed that the expression of 19 genes in cohort1_215_M4, 39 genes in cohort2_75_M3 and 40 genes in cohort3_58_M7 are significantly (log-rank p-value < 0.05) associated with the survival of patients, respectively (Figure S6B; Dataset S7). Among these, the high expression of 19 genes in cohort1_215_M4, 28 genes in cohort2_75_M3, and 31 genes in cohort3_58_M7 are significantly associated with an unfavourable survival of patients. For example, the high expression of *FDFT1* (log-rank p-value = 6.54×10^-4^) with the highest connectivity in both cohort1_215_M4 and cohort2_75_M3 and *MVD* (log-rank p-value = 1.26×10^-3^) with the highest connectivity in cohort3_58_M7 are significantly associated with poor patient outcome (Figure S6C&D). In addition, some of these genes have already been described as associated with NAFLD associated HCC (NAFLD-HCC). For instance, the high expression of *SQLE*, a second rate-limiting enzyme involved in *de novo* cholesterol synthesis with relatively high connectivity in disease-associated modules (Figure S4; Figure S5A), was predictive of unfavourable survival of HCC patients (log-rank p-value = 7.39×10^-4^; Figure S7). Indeed, recent studies have demonstrated that *SQLE* acts as an independent prognostic factor in patients with HCC associated with NAFLD, and *SQLE* inhibition suppressed NAFLD-HCC growth *in vitro* and *in vivo* (Liu et al., 2018a; Ray, 2018).

### Identification of transcription factors that regulate the NAFLD-associated modules

To investigate the transcriptional regulation in maintaining homeostasis and alterations in the disease state, we performed TF enrichment analysis (hypergeometric test) using the genes from the disease-associated modules and module 3 in GTEx, which shows high similarity to disease-associated modules (Figure 5; Dataset S8), based on TRRUST database (Han et al., 2018). Our results indicated that *HNF4A*, *HNF1A*, *PPARGC1A*, *SREBF2* and *PPARA* are the most significantly enriched TFs in GTEx_M3 (Figure 5A). Meanwhile, we observed that *HNF4A*, *SREBF1*, *SREBF2, YY1* and *KLF13* are the significantly enriched TFs in all three disease-associated modules. We also found the significant upregulation of hepatic expression of *SREBF1*, *SREBF2*, *HNF4A*, and *KLF13* in NAFL and NASH compared to the control group (adjusted p-value < 0.05, Figure 5B-F; Dataset S2).

**Figure 5.**
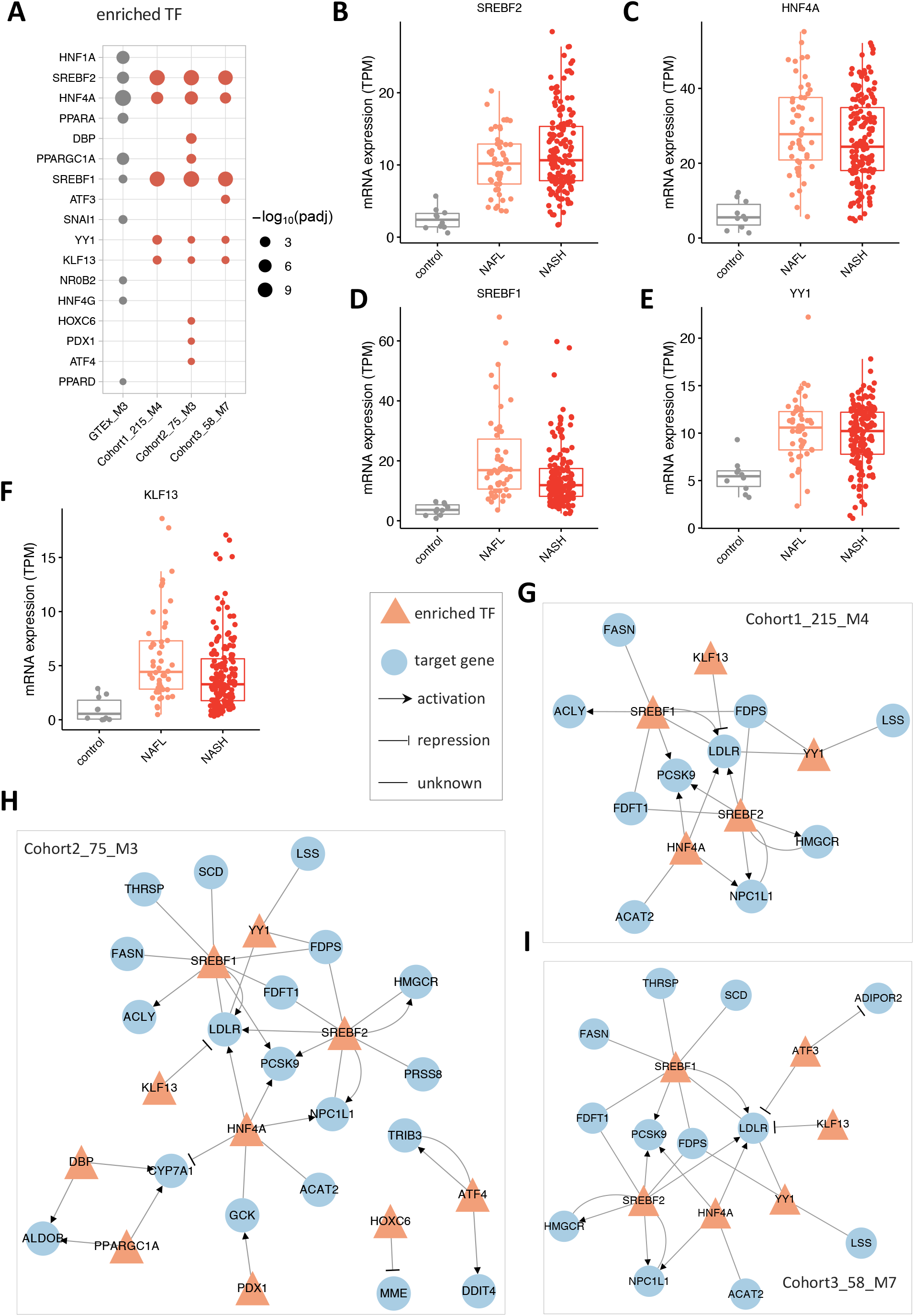
Regulatory relationship between an enriched transcription factor and associated target genes in disease-associated modules. (**A**) enriched transcription factors in GTEx_M3, cohort1_215_M4, cohort2_75_M3, and cohort3_58_M7. (**B-F**) mRNA hepatic expression of the enriched transcription factors in disease-associated modules, including SREBF2, HNF4A, SREBF1, YY1, and KLF13. (**G-I**) the regulatory network between enriched transcription factors and associated target genes in cohort1_215_M4, cohort2_75_M3, and cohort3_58_M7. The regulation between transcription factor and its target was retrieved from the TRRUST database.

We then constructed the regulatory networks for the enriched TFs and associated targets in each of the module (Figure 5G-I; Figure S8). We observed that *HNF4A*, an important transcriptional factor mainly expressed in the liver, regulates the expression of genes involved in lipid metabolism and fatty acid oxidation, including cholesterol/triglyceride transporter (e.g., *ABCG8*, *ABCG5* and *MTTP*), oxidoreductase (e.g., *AKR1C4*, *CYP2D6* and *CYP2B6*) in the regulatory network of GTEx_M3 (Figure S8). As known, *SREBF1* and *SREBF2* regulate the expression of genes associated with *de novo* lipogenesis (DNL) (e.g., *FASN*, *SCD*, *ACACB*), synthesis and cellular uptake of cholesterol (e.g., *HMGCR*, *FDFT1*, *NPC1L*), respectively. Moreover, *PPARA* regulates the expression of genes involved in peroxisomal and mitochondrial β-oxidation, including *ACSL1*, *CPT1A*, *CYP1A1*, and *ACOX1*. Apolipoprotein C3 (*APOC3*), a central regulator of plasma triglyceride levels by inhibiting the removal of remnants of triglyceride-rich lipoproteins, is the most highly regulated gene by *HNF4A*, *NR0B2*, *PPARA* and *PPARGC1A* in the regulatory network of GTEx_M3. Interestingly, low-density lipoprotein receptor (*LDLR*, a key receptor that is internalized by endocytosis) is the most highly regulated genes in the disease-associated modules (Figure 5G-I) by *SREBF1*, *SREBF2*, *HNF4A*, *YY1* and *KLF13*.This indicates that highly co-expressed genes involved in cholesterol metabolism in disease-associated modules are key compared to the other endocytosis-related genes that co-expressed in other modules in the same cohort. In addition to the well-established regulation of *LDLR* activation by transcription factor SREBFs and *HNF4A*, *YY1* and *KLF13*, two specific TFs regulating the disease-associated modules, also showed a regulatory role in the transcriptional regulation *LDLR*. Taken together, the complicated regulation of *LDLR* in the disease-associated modules rather than endocytosis in normal liver tissue might play an essential role in the dysregulation of lipid metabolism underlying the NAFLD pathogenesis.

### Validation of transcription factors in a mouse NAFLD model

Next, we generated liver transcriptomics data from a mouse NAFLD model fed by HSD and performed reporter TF analysis (Huang et al., 2017; Liu et al., 2019; Oliveira et al., 2008) by integrating with the same network of TF-target from TRRUST database (Han *et al*., 2018). We validated the TFs that are enriched in disease-associated modules (Figure 6A). The reporter TF algorithm was used to calculate the statically significant expression changes of gene sets controlled by TFs. To study the regulation of module of genes using this method, we first examined the scored reporter TFs that are significantly associated with the upregulated and downregulated genes in NAFL vs control and NASH vs control, respectively (Dataset S9). The analysis identified 12 reporter TFs of genes in cohort1_215_M4, of which 9 were associated with upregulated genes in NAFLD, including, *ATF4*, *DDIT3*, *HDAC3*, *HNF4A*, *KLF5*, *NFYC*, *SREBF1*, *SREBF2*, and *YY1*. 2 reporter TFs (*VDR* and *WT1*) are associated explicitly with downregulated genes in cohort_215_M4 between patients with NAFL and control samples. *KLF13* was significantly associated with upregulated genes between NASH and control samples. Among these reporter TFs, five of them (*HNF4A*, *SREBF1*, *SREBF2*, *YY1*, and *KLF13*) were also identified by hypergeometric test for cohort1_215_M4 (Figure 5A). Between mice fed by HSD and chow diet (Figure 6A; Dataset S10), 5 reporter TFs (including *KLF5*, *KLF13*, *SREBF1*, *SREBF2*, and *CREB1*) were identified, showing significant association with upregulation of genes using corresponding human orthologs. Taken together, our analysis validated *SREBF1*, *SREBF2*, and *KLF13* as TFs that regulate the hepatic expression of genes in cohort1_215_M4.

**Figure 6.**
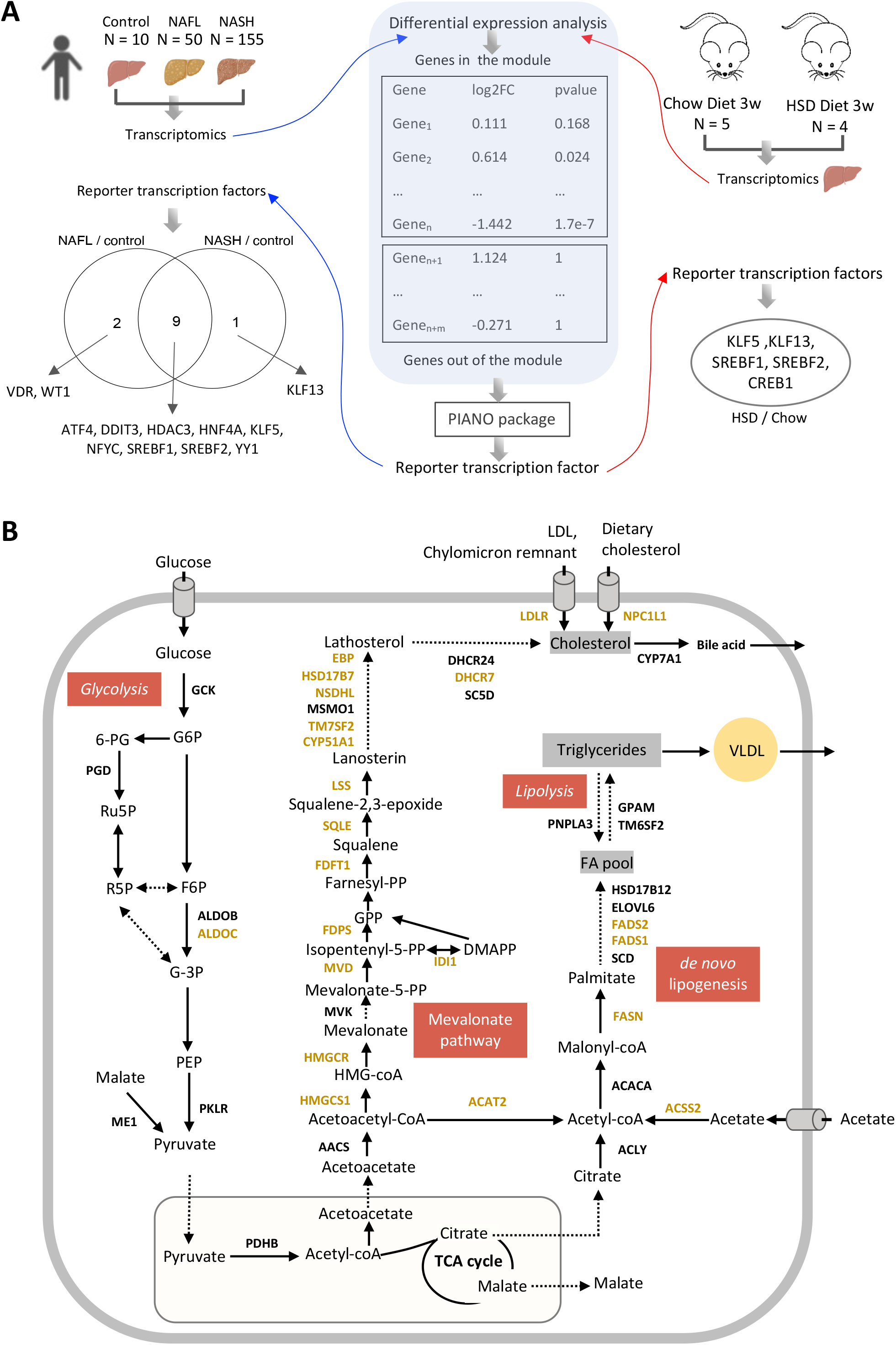
(**A**) Reporter TF analysis was used to validate TFs identified in disease-associated modules using transcriptomics data of NAFLD cohort 1 and newly generated from a mouse NAFLD model. (**B**) Conserved disease-associated modules revealed the dysregulation in the mevalonate pathway, *de novo* lipogenesis, glycolysis, and lipolysis.

### Hepatic co-expression networks reflect dysregulated cholesterol homeostasis and de novo lipogenesis in the NAFLD cohorts

In our network analysis, we found conserved disease-associated modules across three independent NAFLD cohorts, and more than 70% of the genes involved in the modules are associated with metabolic functions. Most of the metabolic genes in this consensus module are associated with cholesterol metabolism. For instance, 13 genes, namely *HMGCS1*, *HMGCR*, *MVD*, *IDI1*, *FDPS*, *FDFT1*, *SQLE*, *LSS*, *CYP51A1*, *TM7SF2*, *NSDHL*, *HSD17B7*, *EBP* and *DHCR7*, which are shared among the three disease-associated modules from different cohorts, and 5 genes, namely *AACS*, *MVK*, *MSMO1*, *DHCR24* and *SC5D* which are included in at least one of the disease-associated modules from different cohorts, are involved in the endogenous synthesis of cholesterol (Figure 6B). *LDLR* and *NPC1L1*, responsible for the uptake of cholesterol, are also found in the disease-associated modules from all three cohorts. In addition, several genes, namely *ACLY*, *ACSS2*, *ACACA*, *FASN*, *SCD*, *FADS1*, *FADS2*, *ELOVL6*, *HSD16B12*, *GPAM*, *PNPLA3* and *TM6SF2*, which are involved in *de novo* lipogenesis and lipolysis, are also included in the disease-related modules in at least one of the cohorts. Finally, a few genes encoding glycolytic enzymes, such as *GCK*, *PGD*, *ALDOB*, *ALDOC*, *PKLR*, *ME1* and *PDHB*, are also found in the disease-related modules. In summary, these indicated a strong connection between the disease-associated clusters with the cholesterol metabolism, *de novo* lipogenesis and glycolysis in the liver and suggested their potential roles in the development of NAFLD.

## DISCUSSION

Here, we applied a systems biology approach on human liver transcriptomics data to elucidate the dysregulated biological processes involved in NAFLD and identified potential regulators via integrating with transcriptional regulatory network. Our analysis identified a highly conserved disease-associated gene module across three different NAFLD cohorts, and this module is specified in the disease networks. At the same time, we could not be found in the network generated from normal subjects (GTEx cohort). Therefore, this gene module could play a critical role in the development of NAFLD and is closely related to the mechanism of the disease. Interestingly, we found the majority of the genes (∼70%) in these disease-associated modules identified in the NAFLD cohort are included in the big gene module 3 of the GTEx cohort, which has 1,975 genes, and this suggested that the disease-associated module and its related biological functions were co-regulated with a large gene group in normal subjects and dysregulated with the progression of NAFLD.

In addition, our analysis provided enriched TFs that regulate the disease-associated modules, which can facilitate our understanding of the regulatory mechanism of these perturbed biological processes. The results from transcription regulatory networks analysis indicated that *SREBF1*, *SREBF2*, *HNF4A*, *YY1*, and *KLF13* are the most prominent regulators of gene expression in disease-associated modules, 3 of which (*SREBF1*, *SREBF2*, and *KLF13*) were validated using the transcriptomics data generated from a mouse NAFLD model. Notably, *KLF13* is reporter TF, specific to this disease-associated module but not for the module from the normal subjects, suggesting their potential role in the development of NAFLD (Ericsson et al., 1999; Natesampillai et al., 2006). It has been shown that selective overexpression of *YY1* resulted in massive triglyceride accumulation and moderate insulin resistance in mice fed with HFD (Lu et al., 2014) and it may a promising target for fatty liver diseases (Wu et al., 2017). We also found that *LDLR* is a central target gene regulated by the enriched TFs in this disease-associated module. It has been demonstrated that multiple mechanisms are involved in protecting against excessive cholesterol accumulation in the liver (Goldstein and Brown, 2009; Natesampillai *et al*., 2006). *LDLR*-mediated endocytosis contributes to this process by removing approximately 70% circulating cholesterol-enriched LDL and providing feedback transcriptional regulation of cholesterol synthesis through SREBFs (Goldstein and Brown, 2009).

Our systematic analyses also highlighted the significant reporter metabolites involving in CS and HS biosynthesis, glycerophospholipid metabolism, folate metabolism and oxidative phosphorylation. Such metabolites are consistent with the findings of previous studies and could be targeted for discovery of potential biomarkers in diagnosis of NAFLD. We also found that most of the genes involved in the disease-associated module are involved in metabolic pathways such as cholesterol metabolism, DNL and glycolysis.

The liver plays a central role in cholesterol homeostasis, and growing evidence has shown that excess hepatic cholesterol causing hepatic lipotoxicity is a predominant factor in the development of human NAFLD (Ioannou, 2016; Min et al., 2012). Abundant hepatic free cholesterol stimulates Kupffer cells and hepatic stellate cells (HSCs), which are key mediators of fibrosis and inflammation as well as mitochondrial dysfunction and thus reflects the severity of disease (Musso et al., 2013). Notably, the differential expression (DE) analysis pointed out significant upregulation of critical genes (adjusted p-value < 0.05) in these cholesterol-related pathways which are involved in the disease-related module, including *HMGCR* (the principal rate-limiting enzyme in mevalonate pathway), *NPC1L1* (a major gene in intestinal cholesterol absorption) in NASH compared to control group (Dataset S2). We found scavenger receptor class B type I (*SCARB1*), which mediate the uptake of HDL cholesterol directly, significantly increased in NAFLD patients compared to the control group. This suggests that upregulation of pathways in both synthesis and absorption of cholesterol may associate with the increased hepatic cholesterol (Ioannou, 2016), as well as increased bile acids in NAFLD patients (Jiao et al., 2018).

Additionally, recent studies have shown that high dietary cholesterol in the mice model is the causative factor for the progression of steatohepatitis to fibrosis and HCC and drives NAFLD associated HCC (Liu *et al*., 2018a; Shen et al., 2020; Zhang et al., 2021). We, therefore, investigated whether the disease-associated modules are predictive of patient outcome using the liver cancer dataset. The results from Kaplan-Meier analysis showed that the high expression of ∼41% genes (12 out of 29 genes identified in all three disease-associated modules) (Figure S6&7) are significantly associated with poor survival of patients, for example, *FDFT1*, *MVD*, *DHCR7*, *SQLE* and *MVD* with high connectivity in these modules. Liu et.al. have shown that targeting *SQLE* can efficiently inhibit the NAFLD-HCC in cellular and animal models (Liu *et al*., 2018a). Considering the characteristics of the co-expression mechanism among genes with similar functions, this integrative network analysis revealed detailed molecules involved in the cholesterol mechanism. It proposed more potential therapeutical targets of effective treatment for preventing NAFLD-to-HCC progression.

Moreover, in the disease-associated module, we also found genes associated with DNL. Generally, it is believed that the triglyceride accumulation in the liver of NAFLD patients is caused by elevation of both DNL and fat uptake (Donnelly et al., 2005; Perdomo et al., 2019). However, we do not find any genes related to fat uptake in these disease-associated modules. In fact, *CD36*, the key free fatty acid transporter, is not significantly changed between NAFLD and control group (Dataset S2). The hepatic expression of *FABP5*, another critical transporter for fat, is significantly decreased in the patients than normal subjects. In addition, most of DNL related genes are significantly up-regulated in the NAFL and NASH patients compared to the control group. Taken together, these suggested that the DNL, rather than free fatty acid uptake, is the source of triglyceride accumulation in NAFLD patients.

Finally, we identified a few key enzymes involved in glycolysis as well as insulin signalling pathway are included in the disease-associated module. For instance, *INSIG*, a key player in the insulin signalling pathway is included in the disease-associated modules among all three cohorts. A recent study has reported that *INSIG* is a central regulator in a negative feedback loop ensuring the balance of lipid desaturation and cholesterol composition and loss of *INSIG1* improves liver damage and would healing NASH progression (Azzu *et al*., 2021). In addition,*GCK*, which is a kinase specific to glucose, is involved in the module of cohort 2, and it is significantly up-regulated in the NAFLD patients compared to the normal subjects (Dataset S2). In our previous study, we have reported that the *GCK* up-regulation is associated elevated insulin resistance in patients and suggested an increased influx from dietary glucose (Lee et al., 2016). Moreover, we also observed that *TKFC*, which is also involved in the module and up- regulated in NAFLD patients in cohort 2. It has been reported that increased dietary fructose uptake could cause NAFLD in both mouse and human patients (Jensen et al., 2018; Loomba et al., 2021). Therefore, these results highlighted the associated between NAFLD and insulin resistance, and suggested the potentially important contribution of dietary glucose, fructose as well as sucrose to development of the disease.

## METHODS AND MATERIALS

### Data retrieving and pre-processing

#### Each dataset was pre-processed independently

NAFLD cohorts. hepatic RNA-seq (raw fastq files) of NAFLD cohort 1 (Govaere *et al*., 2020) and cohort 2 (Hoang *et al*., 2019) were retrieved from European Nucleotide Archive (ENA) database (https://www.ebi.ac.uk/ena/) (Amid et al., 2020) under accession numbers SRP217231 (215 biopsy-proven NAFLD patients) and SRP197353 (78 biopsy-proven NAFLD patients), respectively; Hepatic RNA-seq of NAFLD cohort 3 (Azzu *et al*., 2021) with 58 biopsy-proven NAFLD patients were retrieved from the ArrayExpress data repository (Parkinson et al., 2005) under accession number E-MTAB-9815. Principle component analysis (PCA) was used to detect outlier samples (Figure S1) and three outlier samples in NAFLD cohort 2 were removed based on this analysis. Afterwards, gene abundance in both transcripts per million (TPMs)) and raw count were quantified using the Kallisto (Bray et al., 2016) pipeline based on human genome (ensemble 102 version). We subsequently used DESeq2 R package following a standard protocol (Love et al., 2014) to identify differentially expressed genes (DEGs, adjusted p-value < 0.05) and performed KEGG pathway enrichment using the Platform for Integrative Analysis of Omics (PIANO) R package (Varemo et al., 2013).

GTEx cohort. The RNA-seq data with gene abundance in transcript TPMs from human tissues was retrieved from Genotype tissue expression (GTEx) (https://gtexportal.org/home/datasets) (Consortium, 2013) and remained the samples with available dataset in liver tissue.

### Transcriptomics data from mouse model

Nine C57BL/6J mice were fed a standard mouse chow diet and housed in a 12-h light–dark cycle. From the age of 8 weeks, the mice were then divided into two groups of 5 mice fed with chow diet, 4 mice fed with high-sucrose diet for 3 weeks, respectively. At the age of 11 weeks, all mice are sacrificed and liver necropsy were taken for RNA sequencing. RNA sequencing library were prepared with Illumina RNA-Seq with Poly-A selections. Subsequently, the libraries were sequenced on NovaSeq6000 (NovaSeq Control Software 1.6.0/RNA v3.4.4) with a 2x51 setup using ‘NovaSeqXp’ workflow in ‘S1’ mode flow cell. The Bcl was converted to fastq by bcl2fastq_v2.19.1.403 from CASAVA software suite (Sanger/phred33/Illumina 1.8+ quality scale). The fastq files for mice were then processed as the way that were used in human datasets mentioned above using a standard protocol of Kallisto (Bray *et al*., 2016).

### Construction of co-expression network and analysis

Considering the dramatic increase in size owing to the many gene isoforms and non-coding RNAs (van Dam et al., 2018), we used the “protein-coding genes” for annotation of RNA-seq dataset and then constructed the co-expression network in gene level. For each dataset, we first filtered out lowly-expressed genes based on their median gene expression level (TPMs <1) and constructed co-expression networks by generating gene pairs Spearman correlation ranks within liver tissue, which was performed using “spearmanr” function from *SciPy* (Virtanen et al., 2020) in Python 3.8. Next, we remained gene pairs with significantly (adjusted p value < 0.05) positive correlation (coefficient > 0) on the network and performed module detection analysis using Leiden algorithm (Traag *et al*., 2019), implemented by Python package *leidenalg* (version 0.7.0) with “ModularityVertexPartition” to find the optimal partition. Modules with less than 30 genes were discarded to be able to get significant functional analysis results in the downstream analysis.

To explore the module similarity between different cohorts, we calculated the Jaccard index, which is simply defined as the size of the intersection between two modules divided by the size of the union of the same two modules, and used hypergeometric test to determine whether the genes in one module significantly overlapped with the genes in another module. The overlap was considered as significant when p-value less than 0.05. Topological and node properties of modules were determined using the *NetworkAnalyzer* (Assenov et al., 2008) plugin implemented in *Cytoscape* (version, 3.8.2) (Cline et al., 2007).

### Functional annotation of modules in co-expression network of cohort

KEGG enrichment analysis. we performed functional enrichment analysis for the gene lists of each module of co-expression network using hypergeometric test, which is implemented by the python package *gseapy* (version 0.9.16; https://github.com/zqfang/gseapy), all gene sets of KEGG pathway were obtained from database source of *Enrichr* (Kuleshov et al., 2016).

Disease enrichment analysis. DisGeNet (Pinero et al., 2017) is a platform integrating information of gene-disease association from several public data sources and literature. In our analysis, the lists of diseases enriched by the gene lists in each network module were retrieved from the DisGeNet database using ToppFun of the ToppGene suite (Chen et al., 2009), all gene sets in detected modules were used as background gene sets. Disease terms with Benjamini-Hochberg corrected p value <0.05 were remained and top 20 for each disease-associated module were presented.

Transcription factor enrichment analysis. We retrieved the human Transcriptional Regulatory Relationship Unravelled by Sentence-based Text mining (TRRUST) v2 database (https://www.grnpedia.org/trrust/) (Han *et al*., 2018) and obtained the lists of transcription factor and associated targets, which derived from 7,148 PubMed articles in where small-scale experimental studies of transcriptional regulation were described. In total, 9,395 TF-target regulatory relationships of 795 TFs and 2,493 targets were supplied as database for *Enrichr* (Kuleshov *et al*., 2016), implemented by the python package *gseapy* (version 0.9.16; https://github.com/zqfang/gseapy).

### Reporter metabolite and reporter transcription factor analyses

To investigate the detailed metabolic differences associated with NAFLD, we first performed reporter metabolites analysis (Varemo *et al*., 2013) using the PIANO R package with topological information from liver-specific GEM *iHepatocytes2322* (Mardinoglu *et al*., 2014). Differential expression level of genes (log2-fold change) in each contrast and corresponding significant levels (p value) were used as input.

To validate the enriched transcription factors in disease-associated modules, we also employed the PIANO R package to perform reporter transcription factor (TF) analysis (Huang *et al*., 2017; Varemo *et al*., 2013) in which log2-fold change and p-value of genes, as well as transcriptional regulatory information of TF-target from TRRUST database (Han *et al*., 2018) were used as input. Considering reporter TFs that control the expression of genes in a specific module, we assigned the p-value of genes that are not in that module as 1 in the analysis.

### TCGA data process and Survival analysis

The transcript-expression level profiles (TPM) had been downloaded from Toil (Vivian et al., 2017) under the project ID of TCGA-LIHC. We screened all samples in TCGA-LIHC cohorts and kept 363 donors with both primary tumour solid tissue samples and clinical information. We only extracted tumour samples with identifier “A” for liver hepatocellular carcinoma and subsequently quantified the mRNA expression by Kallisto (Bray *et al*., 2016) based on the GENCODE reference transcriptome (version 23). Genes with an average TPM>1 were reserved for the following analysis. The clinical information was collected from TCGA Pan-Cancer Clinical Data Resource (TCGA-CDR) (Liu et al., 2018b). Samples with a survival time of zero-day were excluded.

Genes were divided into two groups according to TPM values and examined by Kaplan-Meier survival analysis; the survival outcomes were then compared based on log-rank tests. To choose the best TPM cut-offs for grouping, all TPM values from the 20th to 80th percentiles were used to group the patients. Significant differences in the survival outcomes of the groups were examined, and the value with the lowest log-rank P-value is selected. The R package “survival” and graphics “ggplot” was used during the Kaplan-Meier analysis. Genes with log-rank P values less than 0.05 were defined as prognostic genes. In addition, if the group of patients with high expression of a selected prognostic gene has a higher observed event than the expected event, it is an unfavourable prognostic gene; otherwise, it is a favourable prognostic gene. All analysis were applying on R.

## Supporting information

Supplemental index

## Acknowledgments

AM and HY acknowledge support from the PoLiMeR Innovative Training Network (Marie Skłodowska-Curie Grant Agreement No. 812616) which has received funding from the European Union’s Horizon 2020 research and innovation programme. The computations were performed on resources provided by SNIC through Uppsala Multidisciplinary Center for Advanced Computational Science (UPPMAX) under Project sllstore2017024.

