## Supplemental index for "A network-based approach reveals the dysregulated transcriptional regulation in non-alcohol fatty liver disease"

### Supplementary Appendix

**This PDF file includes:**

Captions for Dataset S1 - S10

Supplementary Figures. S1 - S8

#### SUPPLEMENTARY DATASET LEGENDS

**Dataset S1**. Genes in each module of each cohort.

**Dataset S2**. Result of differential expression analysis for NAFLD cohort 1

**Dataset S3**. PIANO results showing enriched KEGG pathways.

**Dataset S4**. Reporter metabolites identified in this study.

**Dataset S5**. Summary of disease-associated analysis from ToppFun.

**Dataset S6**. Topological properties of genes in disease-associated modules.

**Dataset S7**. Kaplan–Meier (KM) analysis for the genes in the disease-associated modules.

**Dataset S8**. Summary of transcription factor enrichment for genes in modules of each cohort.

**Dataset S9**. List of reporter TFs associated with NAFLD.

**Dataset S10**. Result of differential expression analysis for mouse NAFLD model.

#### SUPPLEMENTARY FIGURE LEGEND

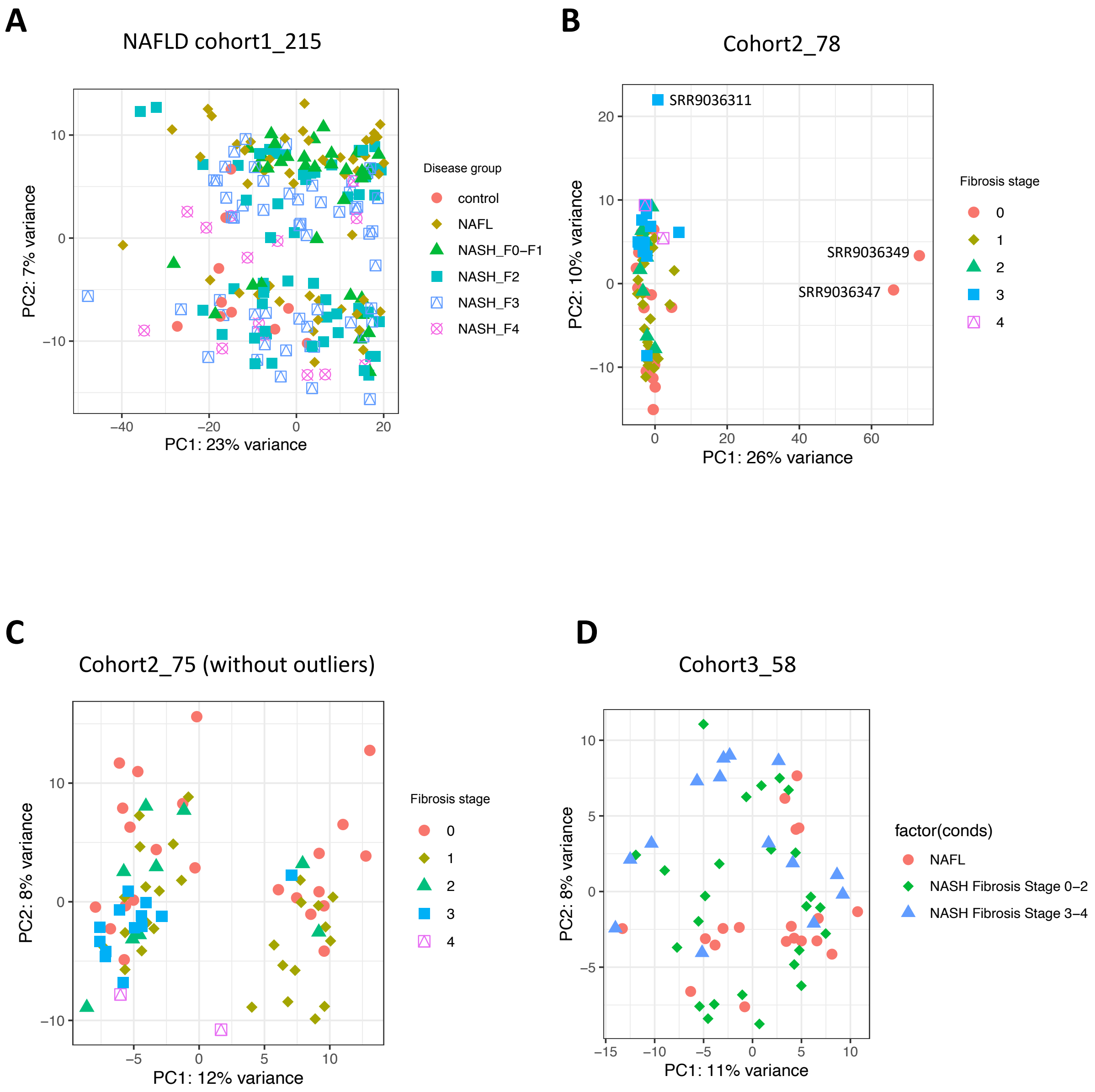


**Figure S1**. PCA analysis for RNA-seq datasets.

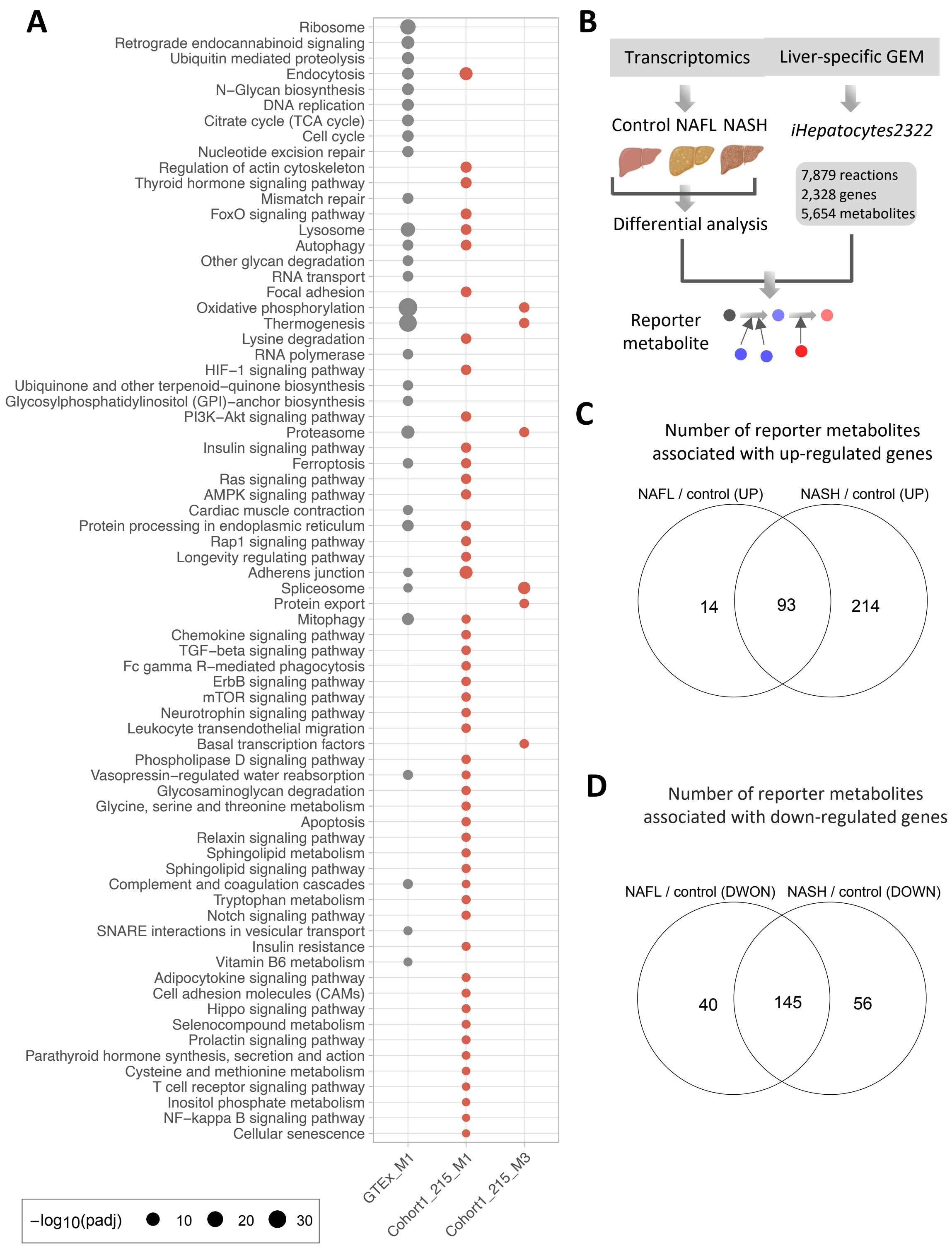


**Figure S2**. (**A**) Dot-plot heatmaps showing KEGG pathways enriched in different modules. Color differences of dots indicate the studied cohort (GTEx or NAFLD) in which the module detected. The size of each dot is proportional to the significance (-log$_{10}$(padj). (**B**) Reporter metabolites analysis was used for the analysis of transcriptomics data from the NAFLD cohort. (**C**) The Venn diagram shows the number of reporter metabolites associated with up-regulated genes in either NAFL vs control or NASH vs control. (**D**) The Venn diagram shows numbers of reporter metabolites associated with down-regulated genes in either NAFL vs control or NASH vs control.

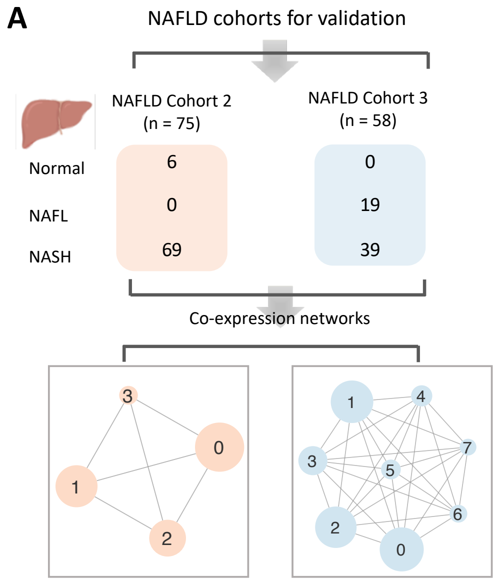


B

| Module size | NAFLD cohort 2 | NAFLD cohort 3 |
|---|---|---|
| Module 0 | 4,009 | 1,548 |
| Module 1 | 3,090 | 1,467 |
| Module 2 | 2,412 | 1,408 |
| Module 3 | 80 | 740 |
| Module 4 | - | 312 |
| Module 5 | - | 236 |
| Module 6 | - | 128 |
| Module 7 | - | 71 |

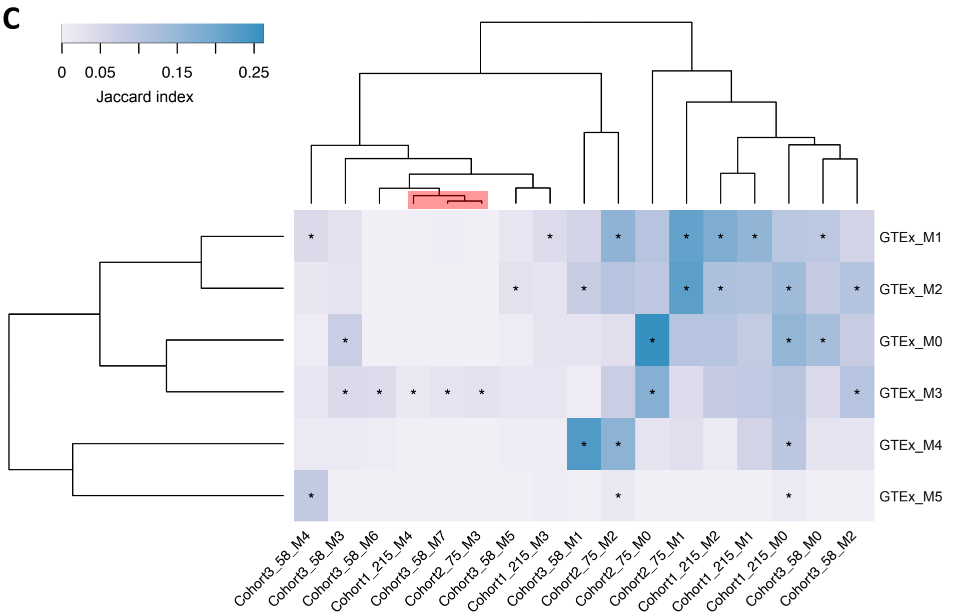


**Figure S3**. (**A**) Sample information of NAFLD cohort 2 and 3 for validation and construction of co-expression networks. Transcriptome data of liver tissue were obtained from NAFLD cohort with 75 and 58 samples ranging from normal, NAFL, NASH, respectively. Spearman rank-order correlation coefficient analysis was applied to calculate the correlation between gene pairs after removing the lowly expressed genes (TPMs <1), and the Leiden algorithm was used to detect modules of significantly correlated genes. The label (number) of the module assigned by the algorithm. (**B**) The numbers of genes consist of the individual module in each cohort. (**C**) Hierarchical clustering of Jaccard Index between module pairs from GTEx cohort and all three NAFLD cohorts. Colour scales representing the range of the Jaccard index. Asterisk indicates the statistical significance of the overlap between gene members in any two modules from the different cohort. The test was performed by a hypergeometric test. The overlap was evaluated as significant when the p-value less than 0.05.

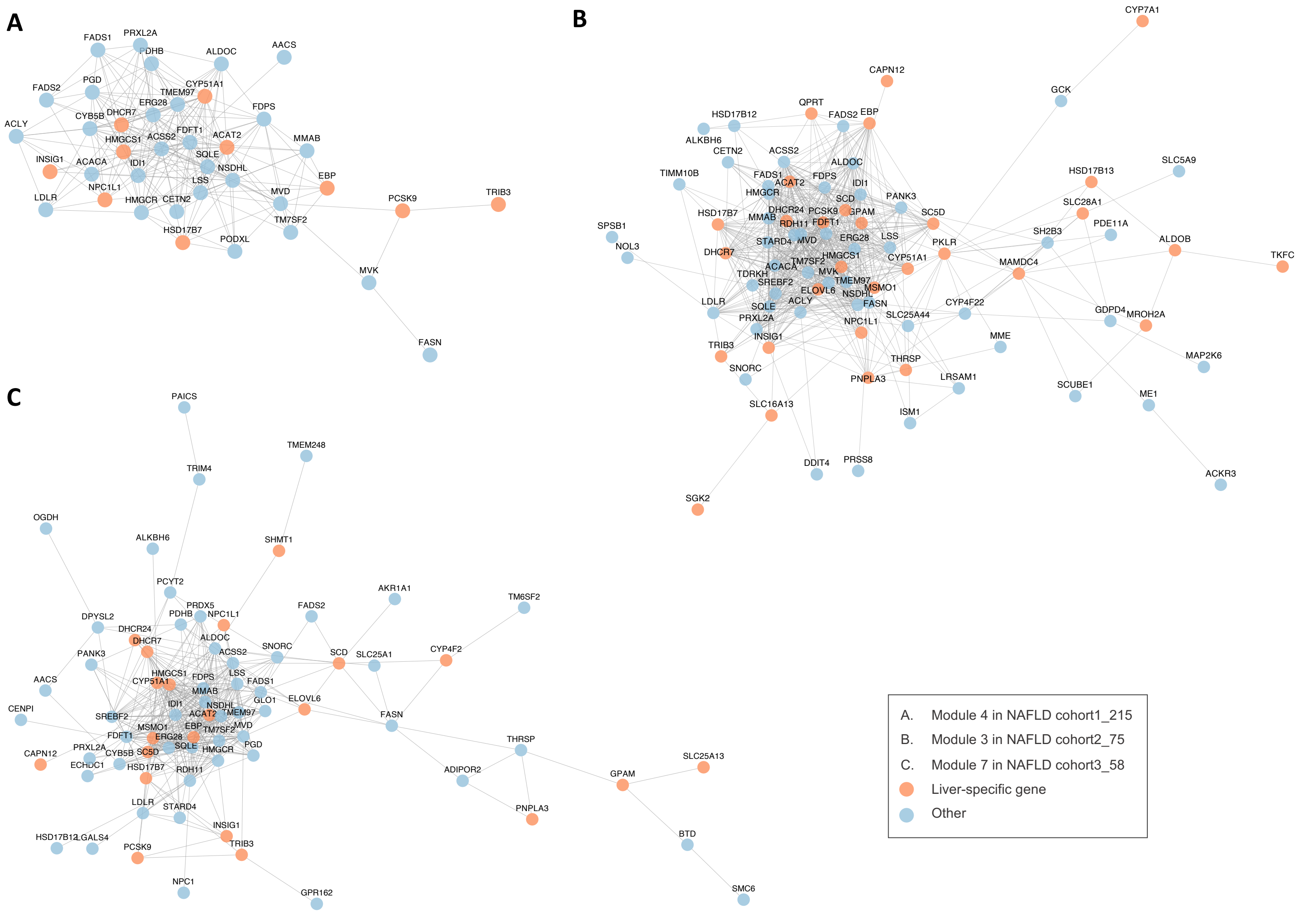


**Figure S4**. Visualization of disease-associated modules constructed by Cytoscape with "Preferred Layout". (**A**) cohort1_215_M4. (**B**) cohort2_75_M3. (**C**) cohort3_58_M7. Orange colour nodes are liver-specific genes based on HPA.

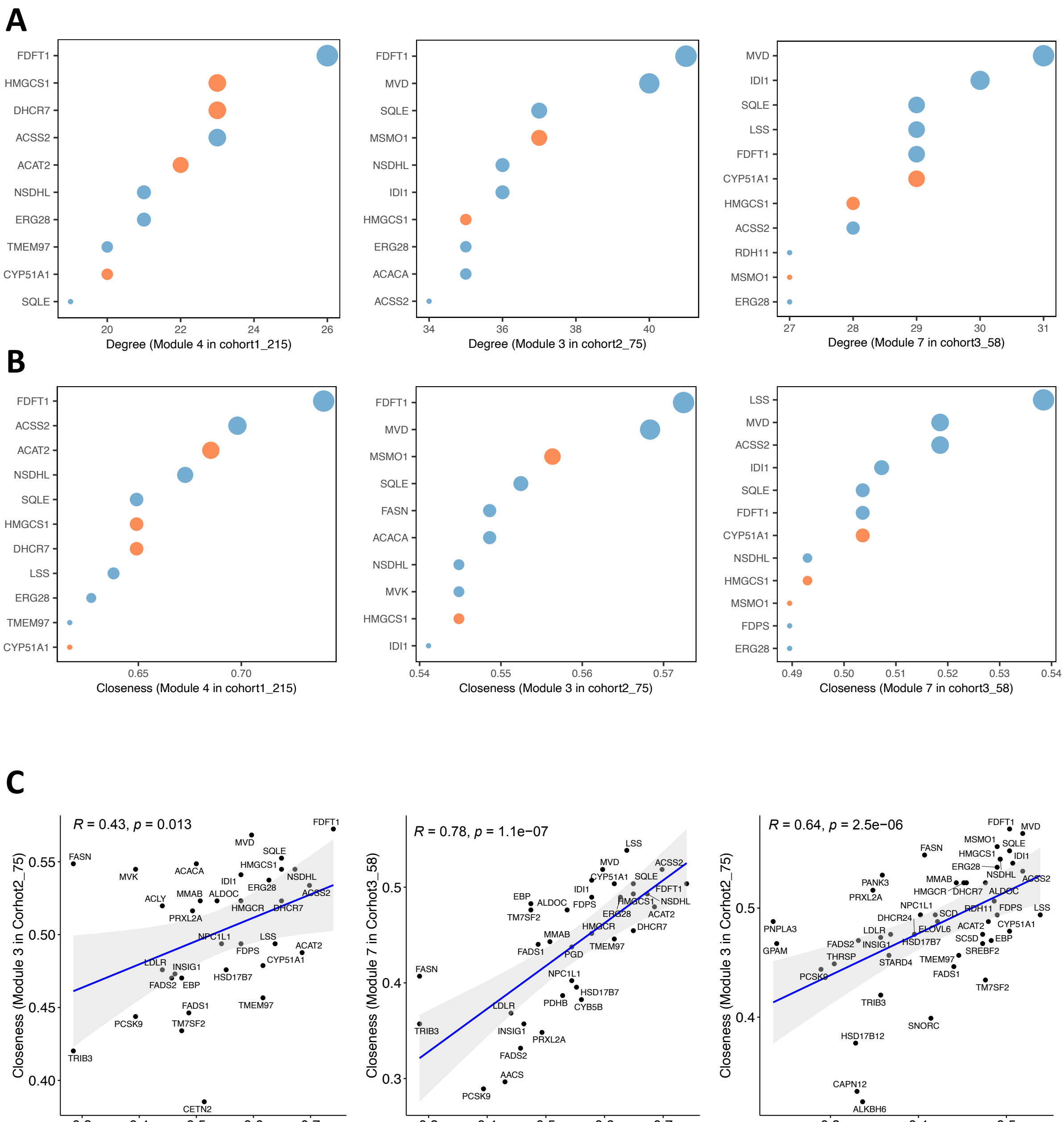


**Figure S5**. (**A, B**) Top 10 genes with highest degree and closeness in disease-associated modules. (**C**) Correlation between closeness among disease-associated modules from different cohorts. The correlation was evaluated by Spearman correlation coefficients.

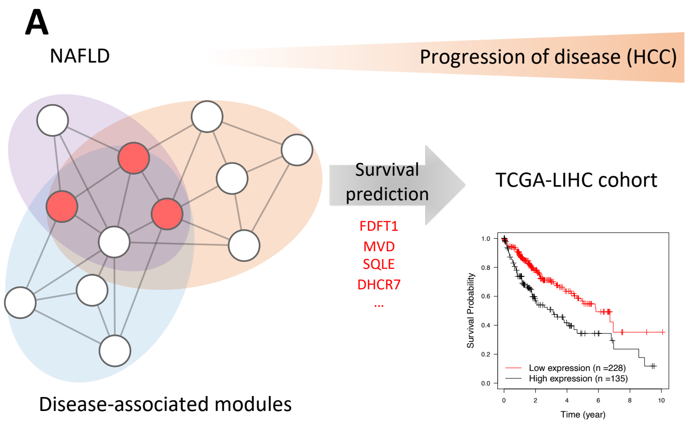


**B**

| Module | Size | KM (Available) | Significant association | favourable survival of patients | unfavourable survival of patients |
|---|---|---|---|---|---|
| Cohort1_215_M4 | 38 | 37 | 19 | 0 | 19 |
| Cohort2_75_M3 | 80 | 77 | 39 | 11 | 28 |
| Cohort3_58_M7 | 71 | 71 | 40 | 9 | 31 |

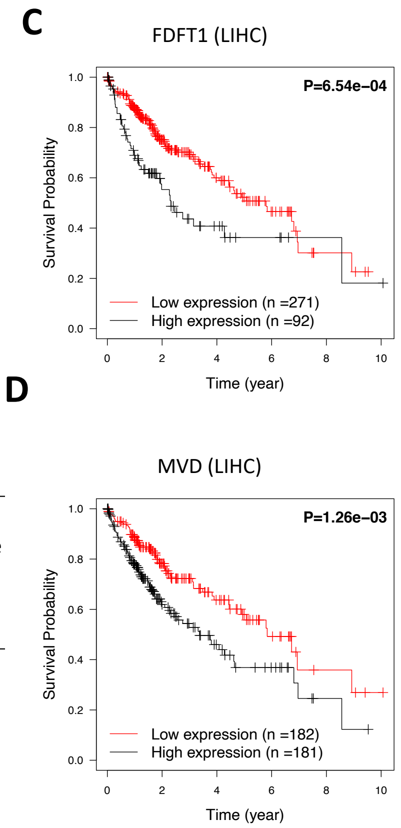


**Figure S6**. (**A**) Schematic framework of survival analysis for the genes in disease-associated modules using the liver cancer dataset in TCGA database. (**B**) Summary of Kaplan–Meier analysis for each of the module. (**C, D**) The Kaplan–Meier plots for FDFT1 and MVD with high-connectivity in cohort1_215_M4 and cohort2_75_M3 and cohort3_58_M7, respectively.

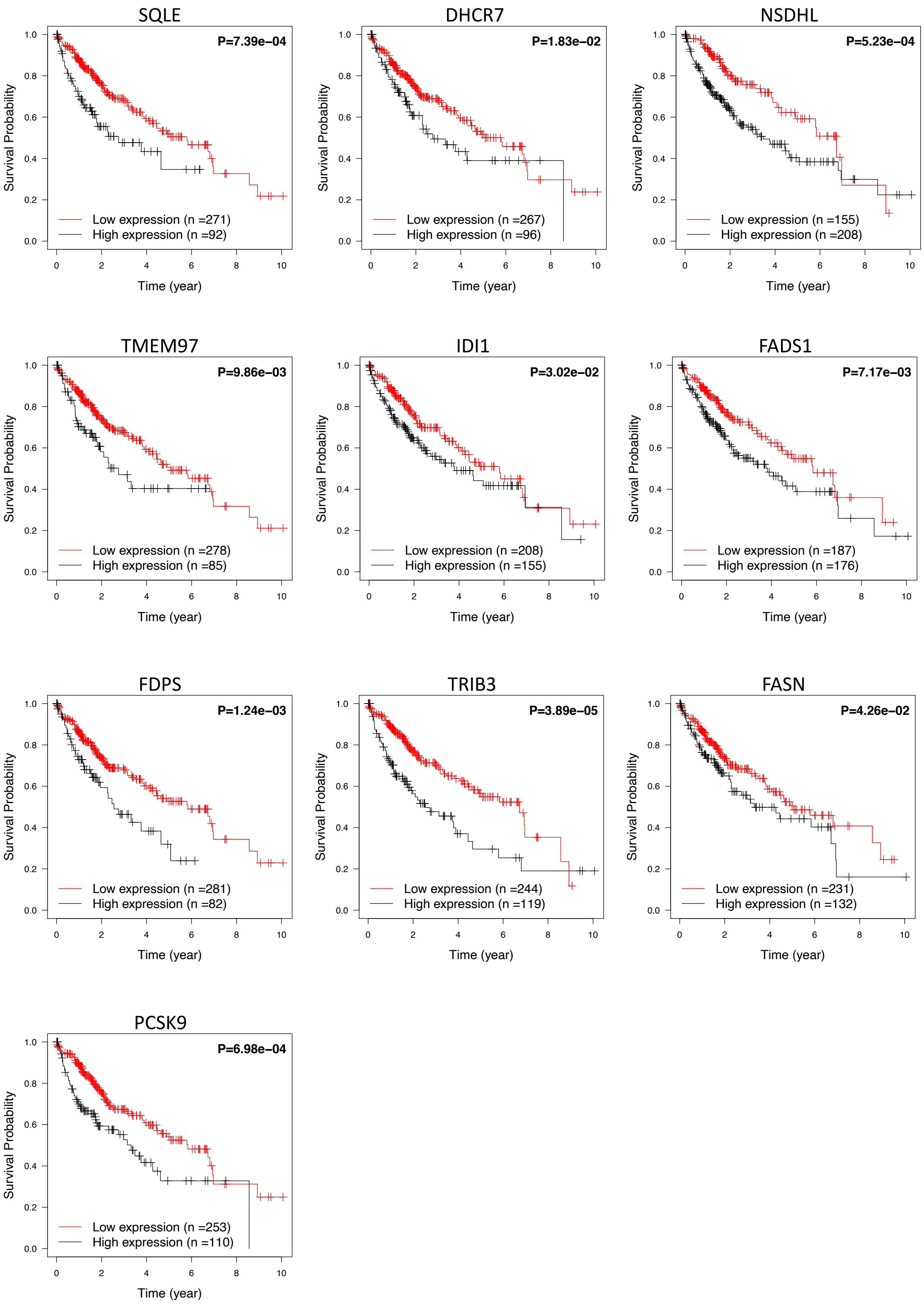


**Figure S7**. The Kaplan-Meier plots showing the high expression of genes shared by disease-associated modules (including SQLE, DHCR7, NSDHL, TMEM97, IDI1, FADS1, FDPS, TRIB3, FASN, and PCSK9) are significantly associated with poor outcome of patients. For each plot, the log-rank test was performed to compare survival curves between high-expression group and low-expression group.

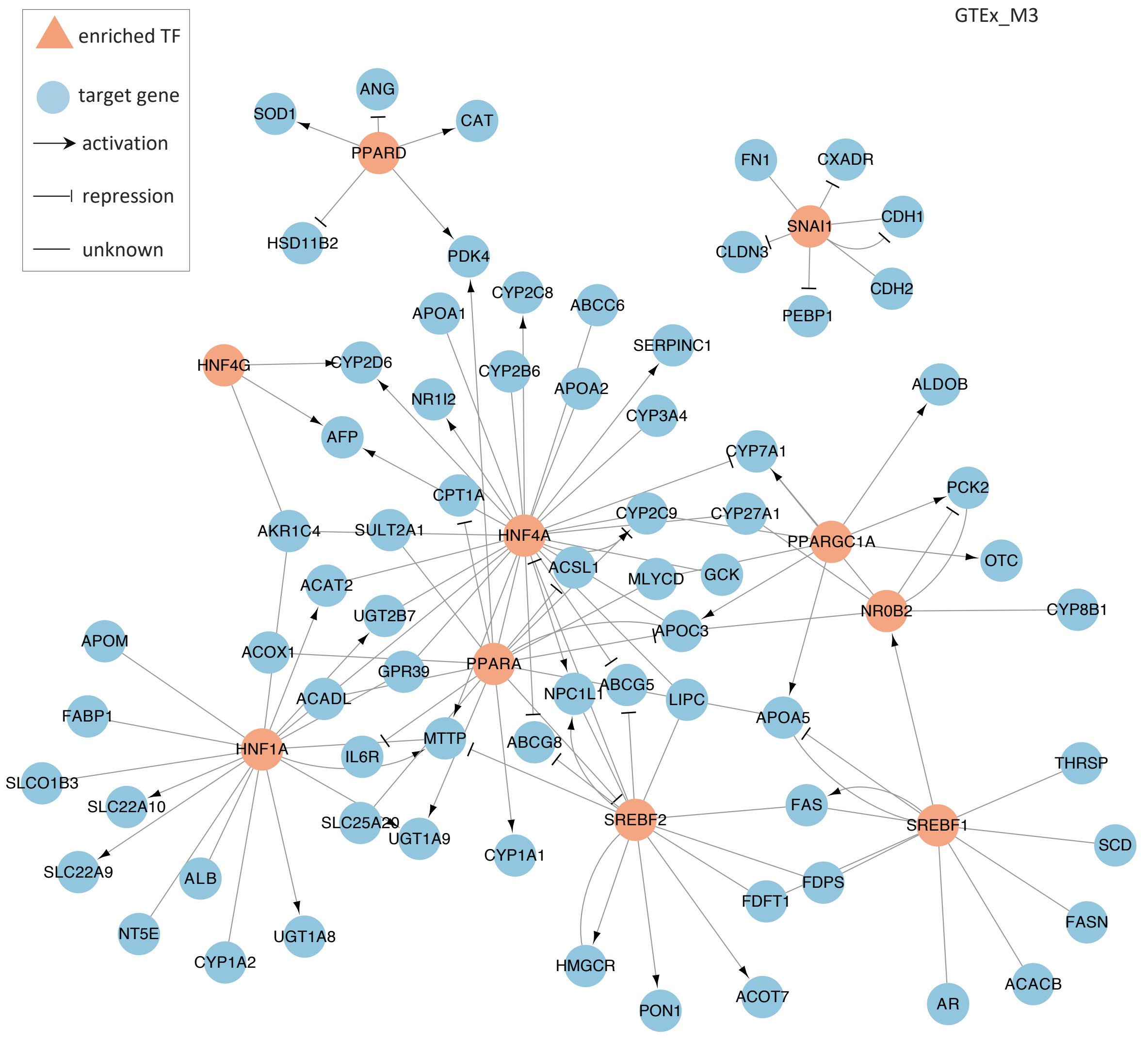


**Figure S8**. Regulatory relationship between an enriched transcription factor and associated target genes in GTEx_M3.
